# “Polar mutagenesis of bacterial transcriptional units using Cas12a”

**DOI:** 10.1101/2020.02.21.959866

**Authors:** Antoine Graffeuil, Bernt Eric Uhlin, David A. Cisneros

## Abstract

Bacterial genes are often organized in functionally related transcriptional units or operons. One such example is the *fimAICDFGH* operon, which codes for type I fimbriae in *Escherichia coli*. We tested the hypothesis that markerless polar mutations could be efficiently engineered using CRISPR/Cas12a in the *fim* operon. Cas12a-mediated engineering of a terminator sequence inside the *fimA* gene occurred with efficiencies between 10 and 30%, whilst other types of mutations, such as a 97 bp deletion, occurred with 100% efficiency. Our results showed that some of the obtained mutants, including one with a single base substitution at the *fim* locus, had decreased mRNA levels of *fimA*, suggesting that the regulation of the *fim* operon was disrupted. We corroborated the polar effect of these mutants by phenotypic assays and quantitative PCR, showing up to a 43 fold decrease in expression of genes downstream *fimA*. We believe this strategy could be useful in engineering the transcriptional shut-down of multiple genes in one single step. For bio-production in *E. coli*, this opens the possibility of inhibiting competing metabolic routes.

## Introduction

The evolutionary success of operons has been attributed to their contribution to the organization of metabolic pathways, which ultimately allowed organisms to become less dependent on exogenous sources of organic compounds (1). Importantly this relationship was maintained during evolution for many metabolic pathways and was probably enhanced by horizontal gene transfer (1). Genes within operons are co-transcribed from a single promoter but it has been shown that the mRNA of each open reading frame (ORF) has an independent folding from their neighbouring genes and this feature directly influences its translation efficiency (2). Other features regulating operon expression and function in *cis* include attenuators, terminators and processive aniterminators (reviewed in (3,4)). All these elements of regulation depend on the formation of secondary RNA structures.

We asked ourselves if sequences with such a tendency to generate secondary structures could be easily engineered in the *E. coli* chromosome using CRISPR/Cas technologies. CRISPR/Cas genome editing in *E. coli* (5), opened the possibility of marker-less genetic engineering with single base resolution. Therefore, it is possible to now generate multi-scale libraries of mutants in a few days (6). The objective of the following study was to design a strategy to shut-down the transcription of entire operons using a synthetic terminator sequence (7) in *E. coli*.

The possibility of easily controlling entire transcriptional units would serve multiple purposes for the bio-production of compounds using synthetic metabolic routes. For example one aspect poorly appreciated in synthetic biology is metabolic burden (8). Protein synthesis has been calculated to be the most expensive step (ATP-wise) involved in cell growth, compared to the biosynthesis of its building blocks (amino acids) which uses only a small fraction of ATP (9). Therefore shutting down the synthesis of unnecessary operons during specific growth conditions could contribute significantly to balance the burden while expressing heterologous operons. Another example is shutting down or the re-direction of competing metabolic routes (10) during bio-production from heterologous operons. To that effect, the bioinformatic analysis of competing or converging metabolic routes will be essential (11). Another interesting application would be the removal of virulence factors and antibiotic resistance genes as a potential antimicrobial strategy (14). These type of genes also tend to cluster together in so called pathogenicity islands. Therefore, the strategy presented in the following study opens another possibility to investigate, with one step, the effect of competing transcriptional units arranged in operons on metabolic burden and compound biosynthesis.

Type one fimbriae are adhesive surface appendages present mainly in *Enterobacteriaceae* that were first shown to agglutinate red-blood cells (15). They are composed of a main pilin subunit repeated in the order of hundreds to thousands and are assembled at the surface by an outer membrane usher. At the tip of the filament the so called minor pilins are assembled in a specific order. The most distal pilin to the cell (FimH) functions as an adhesin (16,17). In the case of *E. coli* type I fimbriae the FimH adhesin is inhibited by D-mannose residues and α-methyl-mannoside and they recognize mannosylated receptors (16–18). Its role in virulence in uropathogenic strains of *E. coli* has been established for a long time (19,20). However, its role in mammalian persistent intestinal colonization has only been recently shown to be essential for the case of uropathogenic *E. coli* (21). It has not yet been studied, how each of the several fimbrial systems present in symbiotic *E. coli* strains (22) contributes to intestinal colonization *in vivo*.

The *fim* operon is a good system to test an engineered polar effect, for several reasons: Firstly, the architecture of the *fim* operon is such that the gene coding for the major and most abundant component of the pilus is the first to be transcribed, followed by the assembly machinery. This gives the advantage that disruption of the first gene inactivates functionality of the fimbriae complex and any polar effect would disrupt the function of the subunits important for fimbriae assembly. Secondly, the levels of *fim* transcription are chiefly determined by an invertible element upstream of the promoter (*fimAp*), which acts as an ON/OFF switch. The regulation of the mannose-dependent adhesion of type I fimbriae is complex, and has been studied *in vitro* for decades. It has been shown to be regulated by various genes including NanR and NagC (trough sialic acid and GlcNAc) (23), Lrp, (24,25), H-NS (26) and IHF (27). All these elements mainly regulate the frequency of inversion of the invertible switch. The molecule (p)ppGpp also regulates the *fim* switch (28). Therefore, it should be possible to quantify a polar effect in the *fim* operon by measuring the transcription of genes downstream of *fimA*.

In this study we introduce a terminator sequence with a strong secondary structure within the *fim* operon to disrupt its transcription. We showed that this sequence is easy to introduce by CRISPR/Cas12a engineering and that the assembly machinery of *E. coli’s* type I fimbriae can be turned off using this polar mutagenesis strategy.

## Results

### Design of *fimA-targeting* crRNA and donor oligos

We set out to disrupt the *fim* operon using Cas12a to introduce a range of different markerless mutations in the *fimA* gene, including the insertion of a terminator sequence (Fig. 1). We designed a CRISPR RNA (crRNA) targeting the *E. coli fimA* gene. To introduce polar and nonpolar mutations in the *fim* operon we designed two complementary oligos for each mutation to form dsDNA donor molecules for homologous recombination. The recombination of these oligos disrupts the binding site of the Cas12a-crRNA complex and permits the positive selection of mutant cells (29).

**Fig 1.**
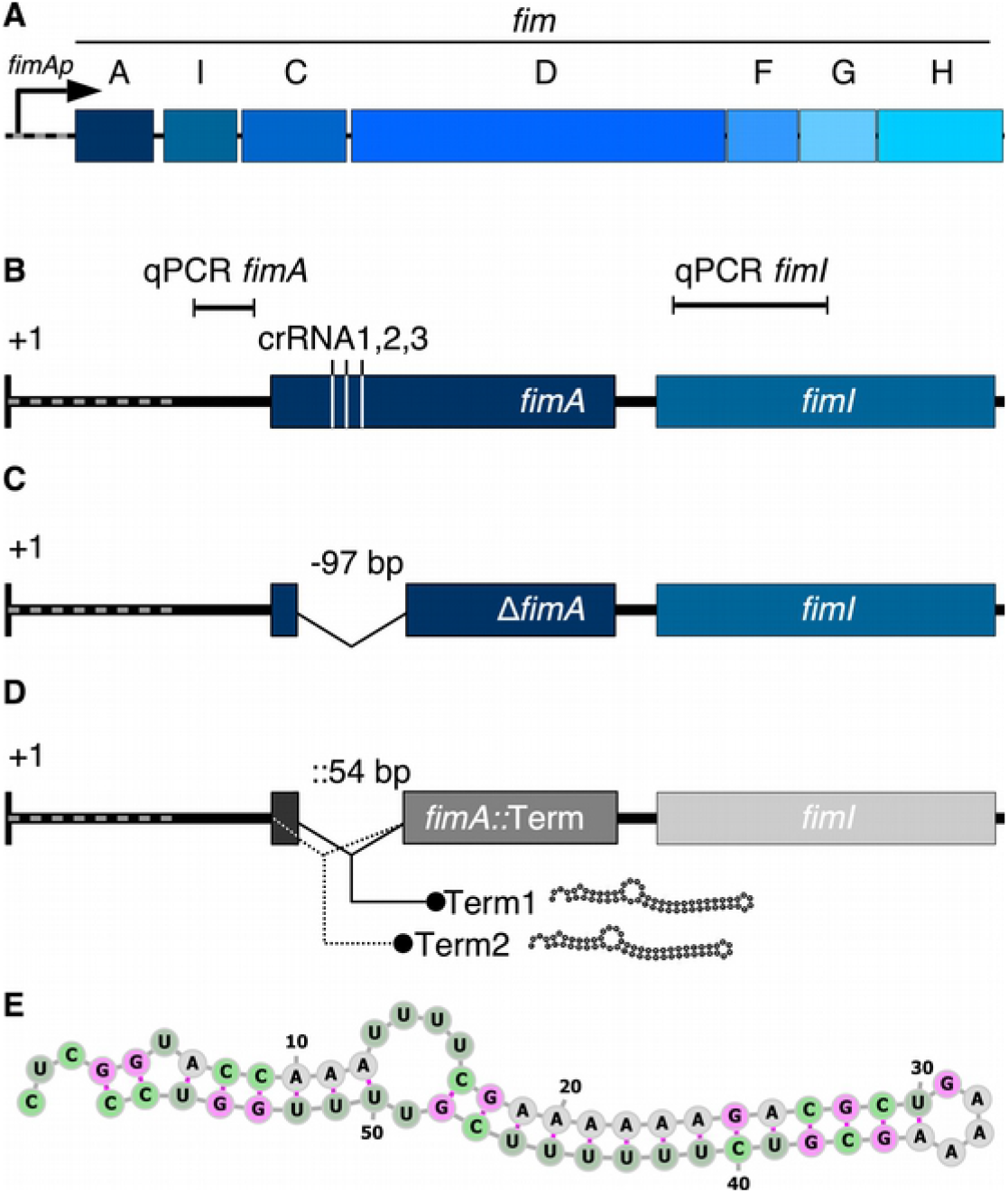
Design of Cas12a-dependent mutagenesis of the *fimAICDFGH* operon. A) The *E. coli fim* operon is transcribed from a promoter (fimAp) containing an invertible element (grey dotted line) and *fimA* codes for the major fimbrial subunit, which was targeted by Cas12a. B) Enhanced view of the *fim* transcript from the +1 transcriptional start site. The regions amplified for qPCR analysis in the *fimA* and *fimI* genes are shown together with the homologous positions to the three crRNAs used in this study. A stop codon was introduced to substitute the PAM sequence to generate a stop codon mutant (*fimA*_*A106T*_). C) Scheme of the 97 bp *fimA* deletion (*ΔfimA*) generated with a dsDNA donor oligo bearing 40 bp of homology. D) Scheme of the 54 bp *fimA* terminator insertion generated with a dsDNA donor oligo bearing a terminator sequence surrounded by 30 bp of homology. Two different insertions were generated: the first was generated at the site of the deletion shown in C (*fimA*_*75*_*∷L3S2P56*) and the second at the position of the start codon (*fimA*_*1*_*∷L3S2P56*) of the *fimA* gene. E) Secondary structure prediction of the L3S2P56 terminator used to create the mutants shown in D.

To introduce a premature stop codon in the *fimA* gene, we designed dsDNA donor oligos in which a single bp change disrupted the PAM sequence adjacent to the target of the designed *fimA* crRNA (crRNA1) and introduced a stop codon (TAA) after 106 bp of the start codon (*fimA*_*A106T*_). To introduce a deletion in the *fimA (ΔfimA)* gene, we designed dsDNA donor oligos containing homology recombination sequences upstream and downstream of the crRNA recognition site to delete 97 bp introducing a stop codon after 75 bp after the start codon (Fig. 1).

To engineer a polar mutagenesis strategy, we designed two different donor oligos introducing a terminator sequence in the *fimA* gene. The first pair of oligos contained the exact same sequences for homologous recombination as the deletion donor oligos but including the L3S2P56 terminator at nucleotide position 75 after the start codon (*fimA*_*75*_*∷L3S2P56*, called here Term1). The second donor oligos were similar but the upstream homologous region corresponded to the leader of the *f imA* gene. Therefore, these oligos introduce a terminator sequence at the start codon (*fimA*_*1*_*∷L3S2P56*, called here Term2).

### Mutagenesis of the *fimA gene*

Terminator sequences have a strong tendency to form secondary structures and therefore we asked ourselves if such sequences could serve as DNA donors for CRISPR/Cas12a mutagenesis. To test this we simply electroporated *E. coli* cells carrying the plasmid pKD46-Cas12a (29) with the annealed dsDNA donor oligos and a plasmid encoding crRNA1. We analysed transformants by colony PCR using oligos that amplify a segment of the *fimA* gene surrounding the site of crRNA targeting. For the stop codon mutant (*fimA*_*A106T*_) obtained with crRNA1, we analysed five colonies. The size of the PCR fragment did not appear to change as compared to wild type (WT) *E. coli* (Fig 2). We purified these PCR products and submitted them for Sanger sequencing. Four of the products showed the specific T->A substitution that introduces a premature stop codon. For the *ΔfimA* mutant, the PCR fragment showed a noticeable change in migration pattern compared to WT (Fig 2). Ten out of ten colonies picked showed the deletion (S1 Fig.). For mutants with the insertion of a terminator sequence only three out of ten and one out of ten colony PCRs showed a change in migration pattern for the Term1 and Term2 mutations, respectively (Fig. 1 and S1 Fig). Altogether these results indicate that the engineering of terminator sequences is possible but with lower success rates compared to the highly efficient engineering of a single base substitution and a deletion.

**Fig. 2.**
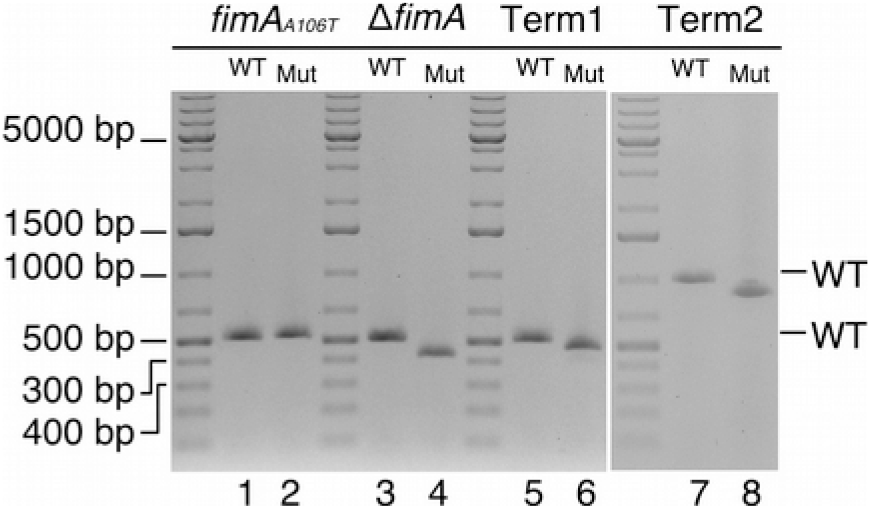
Detection of *fimA* mutants by electrophoresis of PCR products. Lanes 1, 3, 5, show a 514 bp PCR product generated by primers that amplify a segment of the wild type (WT) or mutant (Mut) *fimA* gene. Lane 2 shows the unchanged migration pattern of the PCR product amplifying the stop codon mutant (*fimA*_*A106T*_). Lanes 4 and 6 show the PCR product generated after amplifying DNA from the deletion mutant (*ΔfimA*, 420 bp) and the internal terminator insertion (Ter1, 474 bp), respectively. Lanes 7 show a 988 bp PCR product generated by primers that amplify a segment of wild type *fimA* including the leader sequence and lane 8 shows the 870 bp PCR product generated with DNA from the second terminator insertion (Ter2).

### Phenotypic analysis of *fimA* mutants

To characterise the effect of the different mutations, we verified the functionality of the *fim* operon gene products by a phenotypic assay. We performed agglutination assays of *Saccharomyces cerevisiae* cells (Fig 3A-C). Overnight cultures of WT and *fimA* mutants *E. coli* strains were mixed with a drop of a yeast suspension over a glass plate. With WT *E. coli* this led to agglutination of the yeast cells, which appeared as white, macroscopically visible aggregates (Fig 3D). Contrastingly, mixing *Shigella flexneri* M90T, which is known not to produce fimbrial genes when grown in tryptic soy agar (30), presented no yeast aggregates (Fig 3E). Mixing each of our *fimA* mutants with the yeast suspension produced no agglutination (Fig 3F-I). There was no difference between the stop codon mutant (*fimA*_*A106T*_), deletion and terminator mutants (Ter1 and Ter2), which suggests that the disruption of even one base introducing a stop codon was enough to alter the expression of the *fim* operon.

**Fig 3.**
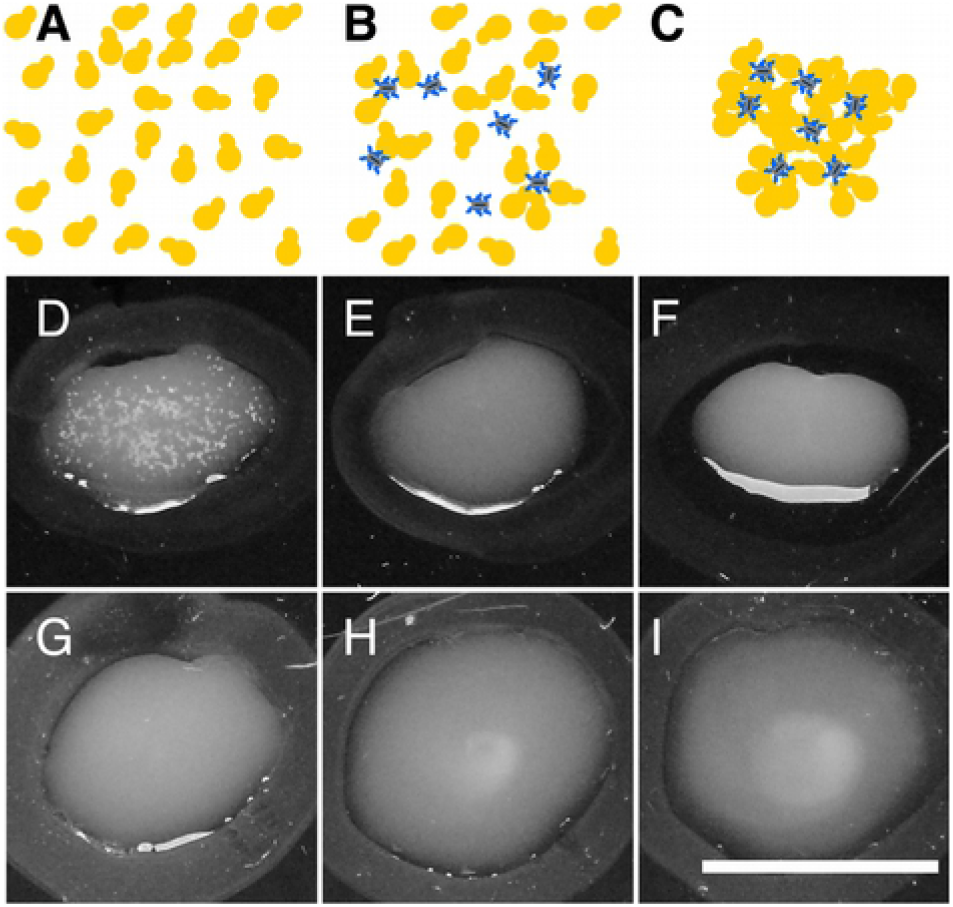
Agglutination assay of *fimA* mutants. A) A suspension of *Saccharomyces cerevisiae* was spread on glass plate. B) The strain to be tested was mixed. The FimH fimbriae subunit recognizes mannose residues on the surface of yeast cells. C) The polyvalent binding of fimbriae to cells causes an agglutination chain reaction, which is macroscopically visible. Agglutination reaction for the following strains: D) WT *E. coli*, E) *Shigella flexneri* M90T (negative control), F) Stop codon mutant (*fimA*_*A106T*_), G) Δ*fimA*, H) *fimA*_*75*_*∷L3S2P56 and* I) *fimA*_*1*_*∷L3S2P56*. Scale bar 2 cm.

### Plasmid trans-complementation of the *fimA* mutants

To verify the efficiency of the different mutations in exerting a polar effect, we verified the functionality of the *fim* operon by a phenotypic assay in WT and mutant strains carrying a plasmid expressing *fimA* (Fig. 4). Each strain was transformed by the trans-complementing plasmid and an empty vector as negative control. WT *E. coli* carrying the empty vector agglutinated *S. cerevisiae* as expected. Similarly, the expression of plasmid encoded *fimA* had no effect on the agglutination reaction (Fig 4A-B), suggesting that neither of the plasmids had an impact on agglutination. Trans-complementation of both the stop codon (*fimA A106T*) and deletion *fimA* mutants resulted in yeast agglutination (Fig 4C-D and Fig 4E-F), suggesting that the mutations did not introduce any effect on genes downstream of *fimA*, which code for the assembly machinery, and that expression of a functional copy of this gene rendered these strains agglutination-positive. In contrast, the strains carrying the empty vector were agglutination-negative. Trans-complementation of both Term1 and Term2 mutants with the empty vector or the plasmid expressing *fimA*, resulted in no yeast agglutination (Fig 4G-H and Fig 4I-J). This suggests that the functionality of the *fim* operon was completely disrupted downstream of *fimA* in these two mutants and suggested that the introduction of the L3S2P56 terminator functionally disrupted the transcription of the entire *fim* operon.

**Fig 4.**
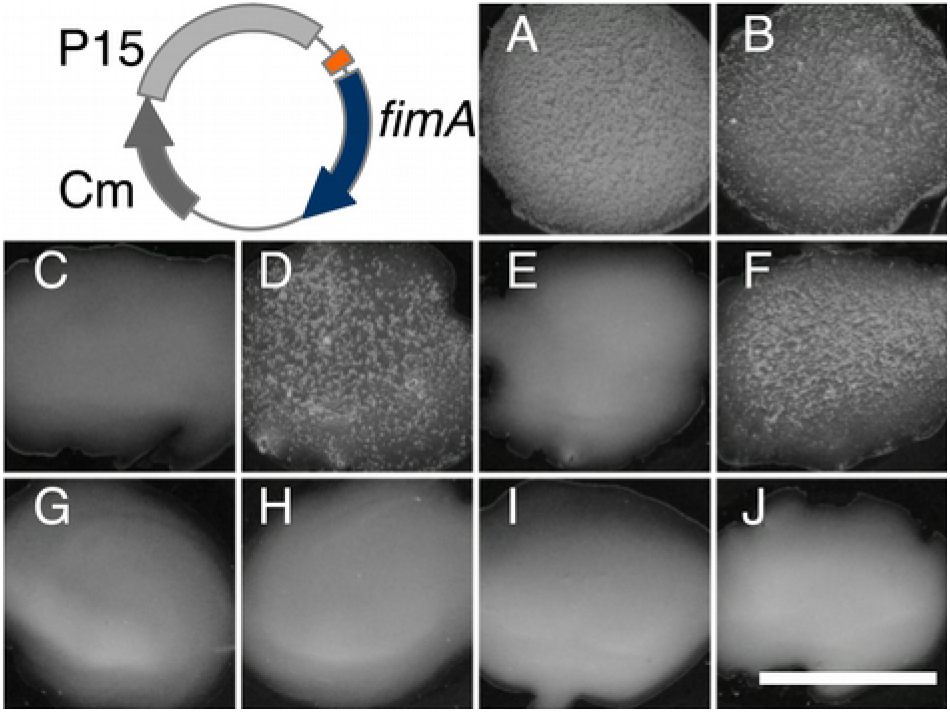
Plasmid complementation of *fimA* mutants. *fimA* was expressed from a plasmid under the *lac* promoter (orange). The empty vector, pSU19, was used as negative control. Agglutination reaction for the following strains: A) WT *E. coli* + empty vector, B) WT *E. coli* + *fimA* plasmid, C) *fimA*_*A106T*_ + empty vector, D) *fimA*_*A106T*_ + *fimA* plasmid, E) Δ*fimA* + empty vector, F) Δ*fimA* + *fimA* plasmid, *G*) Term1 (*fimA*_*75*_*∷L3S2P56*) + empty vector, H) Term1(*fimA*_*75*_*∷L3S2P56*) + *fimA* plasmid, I) Term2 (*fimA*_*1*_*∷L3S2P56*) + empty vector, J) Term2 (*fimA*_*1*_*∷L3S2P56*) + *fimA* plasmid. Scale bar 2 cm.

### Measurement of mRNA levels of *fimA* and *fimI*

We wanted to quantify the extent of a polar effect in all the mutants obtained. We used qPCR to measure the levels of *fimA* and *fimI* mRNA. We designed two sets of primers that allowed for the estimation of the levels of these two genes (Fig 1B). We performed qPCR using as template cDNA derived from mRNA obtained from *E. coli* cultures grown in the same conditions as those used for the phenotypic assays. First, we estimated the amount of the *fimI* mRNA relative to *fimA*. This represents an unbiased measurement that does not depend on the ON/OFF state of the *fimAp*. Relative expression of *fimI* in WT *E. coli* was ~2.5 times lower to *fimA* (Table 1). The median relative expression of *fimI* in the stop and deletion mutants was lower but in the same order of magnitude (Fig 5A). The median relative expression of *fimI* in the Term1 and Term2 mutants was considerably lower (Fig 5A). The relative expression fold change for the Term1 and Term2 mutants was 9 and 43 fold, respectively (Table 1). These results suggest that at least an estimated 9 fold change in expression is sufficient to disrupt the functionality of the *fim* operon downstream of *fimA*.

**Table 1.** Relative levels of *fimI* to *fimA* mRNA. **Relative levels of *fimI* transcript with respect to *fimA***. The median ΔCt values are shown as in **Fig 5A**. The fold change was calculated by dividing the median ΔCt (*fimI – fimA*) of each mutant by the median value in the wild type.

| Strain | Median Δct (*fimI* - *fimA*)* | Median Δct (*fimA* - *fimI*) | ~Fold change inhibition |
| --- | --- | --- | --- |
| WT | 0.39 | 2.6 | 1 |
| Stop codon | 0.10 | 9.8 | 4 |
| Δ*fimA* | 0.17 | 5.8 | 2 |
| Term1 | 0.04 | 23.3 | 9 |
| Term2 | 0.009 | 111.4 | 43 |

**Fig 5.**
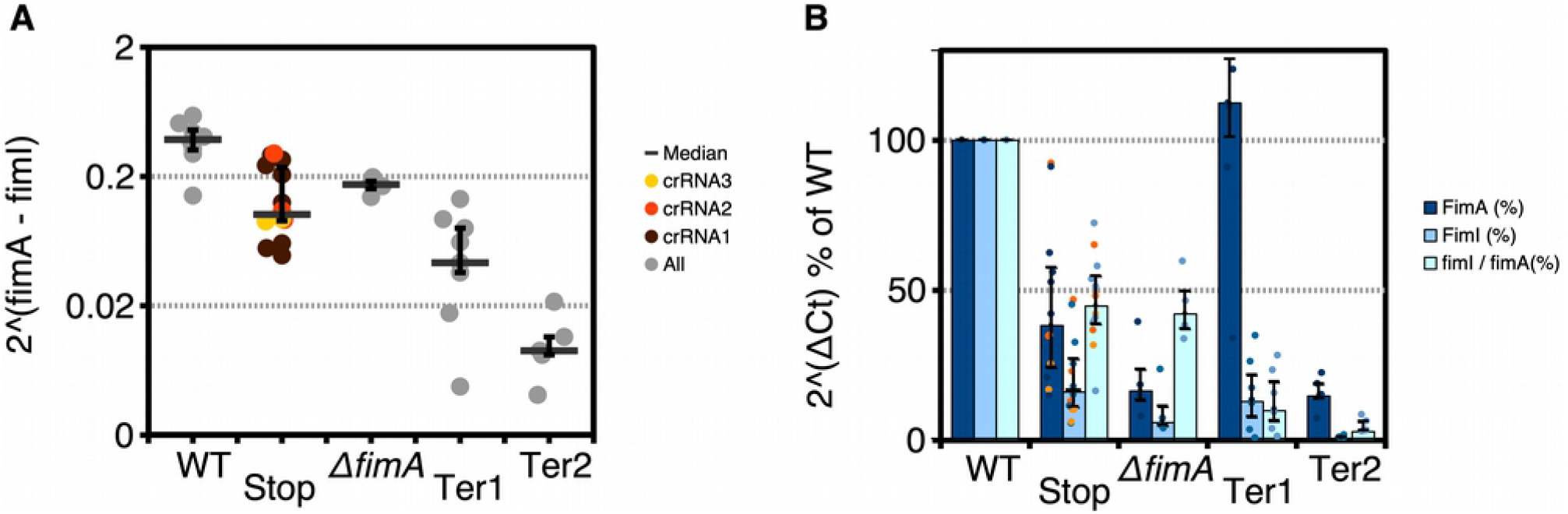
Quantitative PCR analysis of *fimA* and *fimI* expression. Two PCR product were used to quantify the expression of *fimA* and *fimI* (see scheme in Fig. 1). A) *fimI* expression normalized to *fimA* expression. Each data point represents a biological replicate and the median is shown (−). The error bars show the 25th and 75th percentile. B) Same data as in A) but represented as median ΔCt expression of *fimA* and *fimI* normalized to the housekeeping genes *hcaT* and *cysG* and expressed as percent of the wild type. Orange and yellow data points represent the stop codon mutants generated with crRNA1 and crRNA2 as in A. The median ratio of the normalized *fimI/fimA* expression is shown. The error bars show the 25th and 75th percentile. The labels represent the following mutants: Stop (*fimA*_*A106T*_), deletion mutant (Δ*fimA*), Ter1 (*fimA*_*75*_*∷L3S2P56*), Ter2 (*fimA*_*1*_*∷L3S2P56*).

We wanted to measure the levels of *fimI* and *fimA* mRNA levels with respect to housekeeping genes. We designed sets of primers for the housekeeping genes *hcaT* and *cysG*, which have been shown to have stable expression (31). We obtained the average ΔCt with respect to these two genes and normalized it to the WT control performed the day of the experiment. The levels of *fimA* varied in all the mutants as well as the levels of *fimI*. For the stop codon mutant, the deletion mutant and the Ter1 mutant the levels of *fimA* mRNA were lower than the WT (Fig 5B). The levels o f *fimI* were lower for all the mutants, with the Ter1 and Ter2 mutants having the most severe defects compared to WT(Fig 5B).

We were puzzled by the small but measurable changes of mRNA levels in the stop and deletion mutants in *fimA* mRNA(Fig. 5B). To further investigate this phenomenon, we decided to create two different stop codon mutants. We co-introduced in *E. coli* plasmids encoding crRNA2 and crRNA3 (Fig 1) with donor template oligos introducing a stop codon instead of the PAM sequence into *E. coli* carrying the plasmid pKD46-Cas12a. For crRNA2 the donor oligos introduced the mutations T103A, T104A (*fimA*_*T103A,T104A*_) to disrupt the PAM sequence. For crRNA3 the donor oligo introduced the mutations A158T,A159T, A160T,C161A, C162A (*fimA_A158T,A159T,A160T,C161A,C162A_*). Two clones of each of these mutants were selected and analysed by qPCR measurements of *fimI* and *fimA*. The median relative expression of *fimI* to *fimA in* these two new mutants was also lower but in the same order of magnitude of mutants carrying the allele *fimA*_*A106T*_ (Fig 5A, orange and yellow circles). This result suggests that the reduced levels of *fimA* expression are consistent in all three stop codon mutants. Moreover, these results indicate that the lower *fimA* mRNA levels are not caused by an off-target effect of crRNA1.

### Observation of a *fimA* terminator mutant by AFM

We wanted to observe the formation of fimbriae by Atomic Force Microscopy (AFM) and agglutination in previously described favourable conditions. Fluorescence microscopy measurements have shown an increase of fimbriae in static cultures of *E. coli* strain MG1655 (32). We grew WT *E. coli* and the Ter2 mutant expressing *fimA* from a plasmid on static cultures to observe if there was any formation of fimbriae. To prepare the AFM samples in a way that it introduced the least amount of manipulations, we diluted the static culture in deionized water to an OD600 of 0.05 and we immobilized a drop of this suspension on freshly cleaved mica for 20 minutes. After the incubation period the mica was gently washed with more deionized water. In the wild type strain expressing plasmid-encoded *fimA*, we observed bacteria showing the classical type I fimbriae (Fig. 6A and 6A inset). However in the Ter2 mutant expressing plasmid-encoded *fimA* we did not observe any formation of fimbriae (Fig. 6A). In some cells, only the formation of flagella could be observed (Fig.6A inset). To confirm the absence of functional fimbriae in this mutant, we repeated the yeast agglutination assay after static growth conditions (Fig. 6B). As a negative control we incubated this time the wild type strain with α-methyl-mannoside. The wild type strain expressing plasmid-encoded *fimA* was agglutination-positive as shown in Fig. 4. However using the same strain pre-incubated with α-methyl-mannoside completely abolished agglutination. This confirms that agglutination is mediated by the mannose-dependent type I fimbriae encoded in the *fim* operon. Agglutination by the Ter2 mutant expressing plasmid-encoded *fimA* did not complement agglutination after having grown in static conditions and the agglutination reaction appeared as the α-methyl-mannoside negative control. All these results confirm that the Ter2 mutant has a *fim* negative phenotype even when *fimA* is expressed in *trans*.

**Figure 6.**
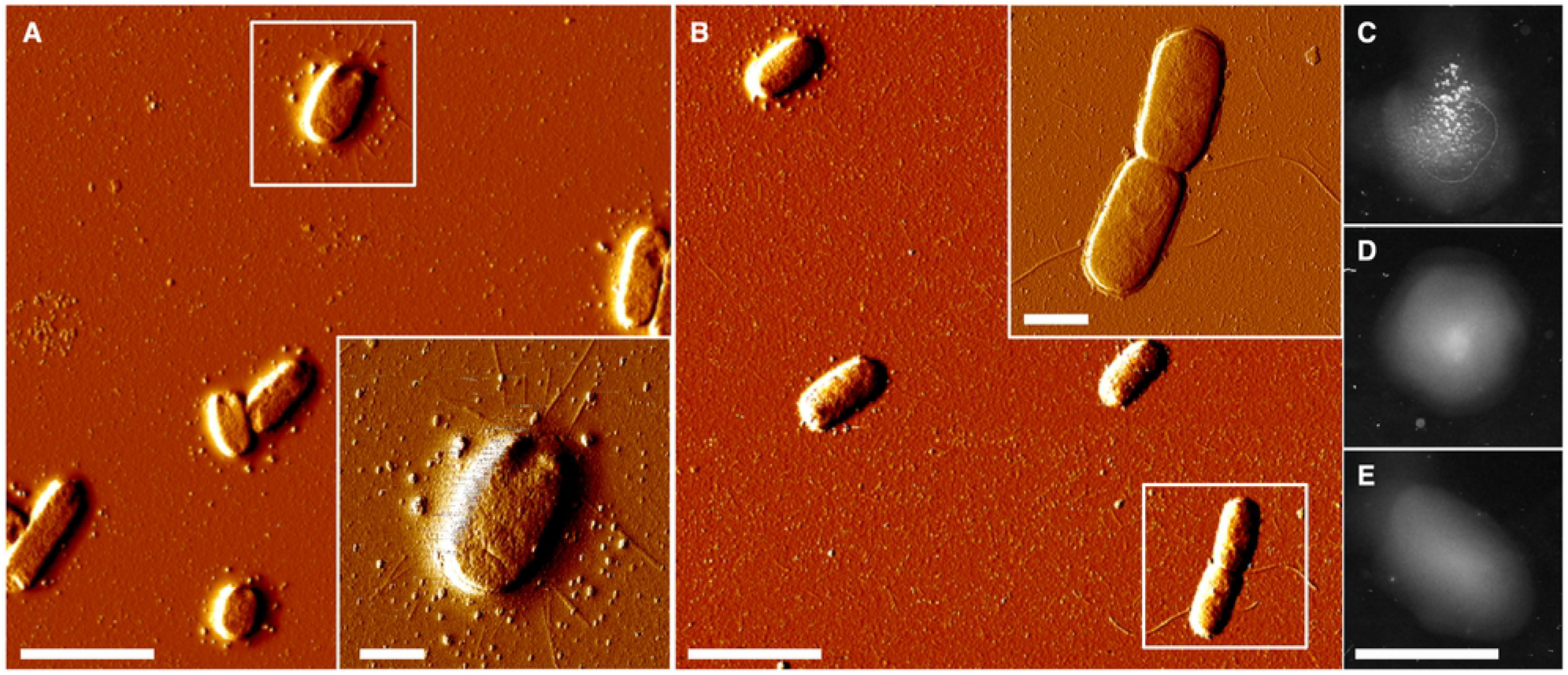
Atomic force microscopy (AFM) and agglutination of static cultures. A) AFM imaging of wild-type *E. coli* + *fimA* plasmid grown in static conditions. A cell showing fimbriae is shown in the inset. B) AFM imaging of Term2 mutant of *E. coli* (*fimA*_*1*_*∷L3S2P56*) + *fimA* plasmid grown in static conditions. A cell with flagella is shown as an inset. C) Yeast agglutination assay of wild-type *E. coli* + *fimA* plasmid grown in static conditions. D) Yeast agglutination assay of of wild-type *E. coli* + *fimA* plasmid grown in static conditions in the presence of α-methyl-mannoside. E) Yeast agglutination assay of Term2 mutant of *E. coli* (*fimA*_*1*_*∷L3S2P56*) + *fimA* plasmid grown in static conditions. Scale bars: A-B) 4 μm. A-B) insets 1 μm. C-E) 2 cm.

## Discussion

CRISPR/Cas technologies have become part of routine work in many laboratories. For the synthetic biology field these techniques opened the possibility of quickly engineer chromosomal loci. For biotechnological purposes, it also opens the possibility to decrease the use of plasmids to produce heterologous enzymes. Plasmids can be lost and impose extra metabolic burdens and regulatory burdens that have to be taken into account when fine-tuning bio-production (reviewed in (8)). One proposed approach is to iteratively test the effect of engineered mutations in strains to optimize this process (33). However the effect of a mutation in a system often carries unpredictable or poorly controlled consequences. In *E. coli*, 63% of its genes are organized in operons (34) and 40% are organised in uber operons (35). In both of these organisation levels, genes tend to be related by functional conservation across bacterial genomes (35). The possibility to transcriptionally shut-down with one step a whole operon could contribute to the quick design and testing of strains with more predictable outcomes if regulons and uber operons were taken into account.

In this study we used an artificial terminator sequence to shut down the transcription of the *fim* operon. This designed artificial sequence has an extended secondary structure that has been shown to improve transcriptional termination (7). We did not know whether such secondary structures could impede the recombination process in the context of the donor homology recombination oligonucleotides used for CRISPR/Cas genome engineering. The predictable outcome from inserting this sequence in the *fim* operon is the premature transcriptional termination of the genes coding for the type I fimbriae assembly machinery. Our results indeed showed a reduction of up to 43 fold in the expression of the *fim* machinery as measured by the expression of *fimI*.

The efficiency of genome engineering strains has been shown to depend on the efficiencies of gRNAs to direct Cas9 DNA cleavage and donor oligos (6). Using the same crRNA (crRNA1) to generate several mutations at the same target sequence we saw that each mutation was produced with efficiencies ranging from 10% to 100%. However the donor oligonucleotides varied in homology arms and secondary structure. Therefore our results suggest that, as suspected, these two parameters are important for bacterial genome engineering. Despite these differences we could readily find mutants with inserted terminator sequences. Among the 582 terminator sequences characterised by Chen *et al.*, we decided to use an artificial terminator sequence (L3S2P56) that showed to have low recombination rates (7), which would ensure more stability within the *E. coli* chromosome. However we did not know whether this would decrease the recombination mediated by homology arms upstream and downstream of the terminator sequence. Our results show that this sequence was easily engineered using the Cas12a-encoding plasmids designed by Yan *et al.* (29). These plasmids are easy to use and allow multiple rounds of mutagenesis similar to previously reported Cas9 approaches (6). Cas12a also brings the advantage that it processes its own crRNAs, which we think will allow certain flexibility in the future when dealing with organisms with poorly characterised promoter sequences. All together this study shows that it is possible to shut down the expression of a whole operon using an artificial terminator. This approach could be complementary to other approaches that allow the transcriptional repression of multiple genes such as the use of multiple gRNAs for transcriptional repression (12) or the deletion of large genomic stretches (13). The use of Cas12a also facilitated genome engineering, because its PAM sequence NTTT (AAAN in the bottom strand) can be easily mutated to introduce a stop codon. Indeed here we showed that a single base substitution (*fimA*_*A106T*_) disrupted PAM recognition and allowed the selection of *fimA* mutant strains.

The *fim* operon was an ideal study case because the phenotype is macroscopically observable and the characterisation of the *fimA* mutants by yeast agglutination produced easy to interpret results. Moreover, the extent of fimbriation is directly correlated to the state of the switch inside *fimAp* (32), therefore for every *fimA* transcript observed the presence of a *fimI* transcript reflects the polarity of the mutants generated here. Indeed, insertion of a terminator sequence abolished the trans-complementation by a plasmid-encoded *fimA* gene. Less predictable phenotypes were obtained by measuring the relative expression of *fimI* to *fimA* in typical laboratory conditions. Three out the four mutants that were generated in this study, both polar and nonpolar, had reduced median levels of *fimA transcript* and of the ratio *fimI/fimA* transcripts. This includes a single base substitution that introduces a stop codon *(fimA_A106T_)*. Several possible explanations exist to accommodate this fact. For example, it is known that ribosome binding protects mRNAs from RNase E degradation (36). Therefore, one possible explanation would be that the premature stop codon introduced in *fimA*_*A106T*_ and in the deletion mutant induces the premature release of ribosome (37,38) and at the same time exposes the *fimA* transcript to degradation. Another more complicated involves the stalling of ribosomes during translation, which may induce mRNA cleavage next to or at a stop codon (39). An initial cleavage event could lead to further degradation by classical mRNA decay pathways. However this could only make sense if the folding of the *f im A* mRNA would be compromised and stalling would be induced by the mutations designed here. This can also be put in the context of the recent discovery that the mRNA of individual genes within operons have a unique secondary structure (2). Deletions smaller than a whole gene could affect the folding of an individual genes’ mRNA within an operon. Another possibility would be that there would be some kind of positive feedback on type I fimbriae expression or functionality. On the conditions that we assayed *fimA* expression, that is rich medium at 37 C, switching off dominates on *fimAp* (40). Therefore, any kind of feedback would quickly result in *fimA* expression inhibition. Another possibility would be that the mutation introduced in this study interferes with a differential stability within *fim* genes’ transcripts, which is an important element for other fimbrial systems (e.g, the *pap* fimbrial operon from uropathogenic *E. coli*) (41). One final possibility is that expression of a functional type I fimbriae system confers a growth advantage (32,42), although this has only been proposed for static cultures. Alternatively, the peptide products of the stop codon mutants (~35 amino acids long and hydrophobic) could also be partially toxic and induce the selection of cells with the *fim* switch in the off state. Although this would not apply to the terminator insertion at position 1 (Ter2) mutant, for which there should not be any partial *fimA* peptide product. Further studies will be needed to clarify the role of the three stop codon mutants and deletion on the expression of *fimA*. However, our results show that even a single nucleotide substitution can have subtle but measurable changes at the transcriptional level within an operon when measured in typical laboratory conditions.

The chromosome of *E. coli* encodes multiple fimbrial systems. However, in the conditions used in this study, only the *fim* type I fimbriae are functional (22). The results shown in this study open the possibility of quickly generating *E. coli* strains in which each fimbrial operon is inactivated. This could be useful to see if any of these systems is essential under a particular growth condition. Another future direction of interest is to characterise the effect of other *fimA* stop codon mutants on the stability of its transcript. It would be crucial to determine if the effects observed here are gene specific or can occur in many other premature stop codon mutants. This is important in the light that introducing a premature stop codon is the easiest form to disrupt a gene using markerless techniques such as Cas9- or Cas12a-mediated mutagenesis.

In summary, in this study we show that Cas12a can be used to introduce markerless insertions in *E. coli* in one step and that this insertion can have strong secondary structures such as terminator sequences without excessively compromising the efficiency of the process. This approach allows the transcriptional shut-down of whole operons as demonstrated for the case of the *fim* type I fimbriae system. More importantly, this strategy can be used to easily test the importance of entire operons for the physiology of *E. coli* or its effect in the design of novel or artificial biosynthetic pathways.

## Materials and Methods

### Cas12a Mutagenesis

To design Cas12a crRNA targeting *fimA* with no off-targets in *E. coli* MG1655 (43), we used the webtool Chop-Chop (44) and crRNAs ranking 1, 2 and 4 were selected due to its closeness to the start codon. To introduce all the mutations designed in this study we used the plasmids designed by Yan *et al.* (29) and followed their protocol with some modifications. Cloning of crRNAs was done by annealing forward and reverse oligos at 95 °C and ramping the temperature down at 2.55°C/min. The oligos were then phosphorylated using polynucleotide kinase (ThermoFisher). The plasmid pAC-crRNAred (29) was digested with BsaI-HF (New England Biolabs), dephosphorylated with FastAP (ThermoFisher) and ligated to the annealed oligos. Donor oligos were treated in the same way as crRNA oligos and their annealing was verified as suggested by König *et al.* (45). The night before the experiment we grew *E. coli* MG1655 carrying plasmid pKD46-Cas12a at 30 °C starting from a glycerol stock. The day of the experiment, 100 μl of overnight culture were transferred to a new agar plate with 0.2% arabinose and 4 h later electrocompetent cells were prepared using the “rapid” protocol (46). These cells were co-transformed by plasmids carrying one crRNA and an annealed pair of donor oligos. As a control transformation we used a crRNA targeting the gene coding for GFP. Colony PCR was then performed on transformants for genotyping. Each colony was resuspended in 20 μl water and boiled for 5 min. One μl of this sample was used as template for PCR. Agarose gel electrophoresis or Sanger sequencing was followed to determine the number of mutants. At least two independent colonies of each mutant were selected for further analysis. After mutagenesis was confirmed by sequencing, the strains were cured from the plasmids following the recommendations of Yan *et al.* (29). We observed that this process had a 100% efficiency by growing the cells at 37 °C in 5% sucrose in very rich medium such as YT2X. Glycerol stocks were then prepared for each clone. The structure of the L3S2P56 terminator was calculated with the Vienna websuite (47) and visualised with Forna (48).

### Agglutination assay

To observe yeast agglutination, wild type or mutant MG1655 was grown overnight for 16 h on Luria agar plates as described before (49). A 1 ul plastic loop was used to take a colony from the plate and mixed with 200 μl of a *Saccharomyces cerevisiae* suspension at OD_600_ 5 prepared from refrigerated baker’s yeast (Lidl Sverige). The loop was used to gently mix the two cell populations and the appearance of white flocculated aggregates could be observed for the wild type strain in less than 30 seconds. *Shigella flexneri* 5A M90T *was* used as a negative control (50). To perform this assay from static cultures, 10 μl from strains grown overnight in static broth (20 ml tryptic soy broth) were mixed with 200 μl of a *Saccharomyces cerevisiae* suspension at OD_600_ 5. In the case of the wild type strain expressing plasmid-encoded *fimA*, the agglutination took approximately 2-5 min under this conditions.. As a negative control, 10 μl of a 3% (w/v) solution of α-methyl-mannoside was added to a 10 μl suspension of wild type *E. coli* and pre incubated for 1 minute before agglutination.

### RNA isolation

To isolate RNA we used the modified method described by Blomberg *et al.* (51) with one further modification. Namely, we grew cells in the same conditions as for agglutination assays (overnight on Luria agar plates) on a lawn. The equivalent to half a plate was scraped and dissolved immediately in DEPC treated water with 10% SDS, 20 mM sodium acetate pH 4.8 and 10 mM EDTA and flash frozen in liquid nitrogen. Frozen cells were thawed and lysed by incubating the samples for 5 min at 65 ºC. Total RNA was extracted by the hot-phenol method (51). Residual DNA was digested by DNase I (ThermoFisher; 1 U/μg RNA, 60 min, 37 ºC) in the presence of RNase inhibitor (Ribolock, ThermoFisher Scientific; 0.1 U/μl) followed by a cleaning step using phenol/chloroform/isoamyl alcohol (25:24:1) and precipitation with ethanol, 0.1 M sodium acetate pH 5.5 and 20 μg of glycogen (ThermoFisher). Removal of residual DNA was verified by PCR using *fimA* oligos. Complementary DNAs (cDNAs) were prepared from each RNA sample using the RevertAid kit (ThermoFisher) according to the manufacturer instructions. As a control of DNA contamination we prepare for each sample a sample without reverse transcriptase (-RT sample).

### qPCR analysis

To perform qPCR we used SYBR Green master mix (ThermoFisher) according to the manufacturer instructions. To analyse the expression of *fimA*, *fimI*, *hcaT* a n d *cysG*, we designed specific oligos using primer BLAST (Table 2). We used an iCycler iQ5 (Bio-Rad) and for each gene cDNAs generated from bacteria grown on plates and as a control we produced a sample without reverse transcriptase (−RT) using the exact same RNA. We verified that for each qPCR experiment we did not find any detectable signal in the -RT sample. Denaturation curves were also acquired to verify the presence of a single product. We performed these controls for every single biological replicate. To verify the linearity of each primer we diluted the cDNA obtained from wild type *E. coli*, which always contained the highest amounts of *fimA and fimI* and verified that 10 times dilutions steps had a slope close to −3.3. To analyse qPCR data we used the iQ5 optical system software (Bio-Rad) obtained the ΔCt from the suggested values of the software. We verified in any case manually that changes in the threshold did not affect or bias the results. Each measurement was quantified using the following formula: Δct=2^(ct1-ct2), where ct is the threshold crossing value at which the SYBR green fluorescent signal appears. In the case of the *fimI* to *fimA* ratio reported in Fig. 5A we simply report the ΔCt without any transformation. Although Ct values varied across experiments, the *fimI* to *fimA* ΔCt was, in our hands, consistent across experiments within one order of magnitude. To normalize the data of *fimA* and *fimI* against housekeeping genes we obtained the ΔCt of each sample against the ct value of *cysG* and *hcaT* and the average of the two values was obtained. The average ΔCt values of each sample was normalized against the ΔCt of the wild type strain as a percentage. The ratio of the *fimI* to *fimA* expression was obtained by dividing their normalized ΔCt values and expressed as a percentage.

**Table 2.** Strains used in this study.

| Name | Strain | Reference |
| --- | --- | --- |
| Wild type | *Escherichia coli* MG1655 | Guyer et al. |
| *S. flexneri* | *Shigella flexneri* 5A M90T | Sansonetti et al. |
| Stop1 | *Escherichia coli* fimA<sub>106T</sub> | This study |
| Stop2 | *Escherichia coli* fimA<sub>T103A,T104A</sub> | This study |
| Stop3 | *Escherichia coli* fimA<sub>A158T,A159T,A160T,C161A,C162A</sub> | This study |
| fimA | *Escherichia coli* fimA<sub>76-172</sub> | This study |
| Term1 | *Escherichia coli* fimA<sub>75</sub>::L3S2P56 | This study |
| Term2 | *Escherichia coli* fimA<sub>1</sub>::L3S2P56 | This study |

**Table 3.**
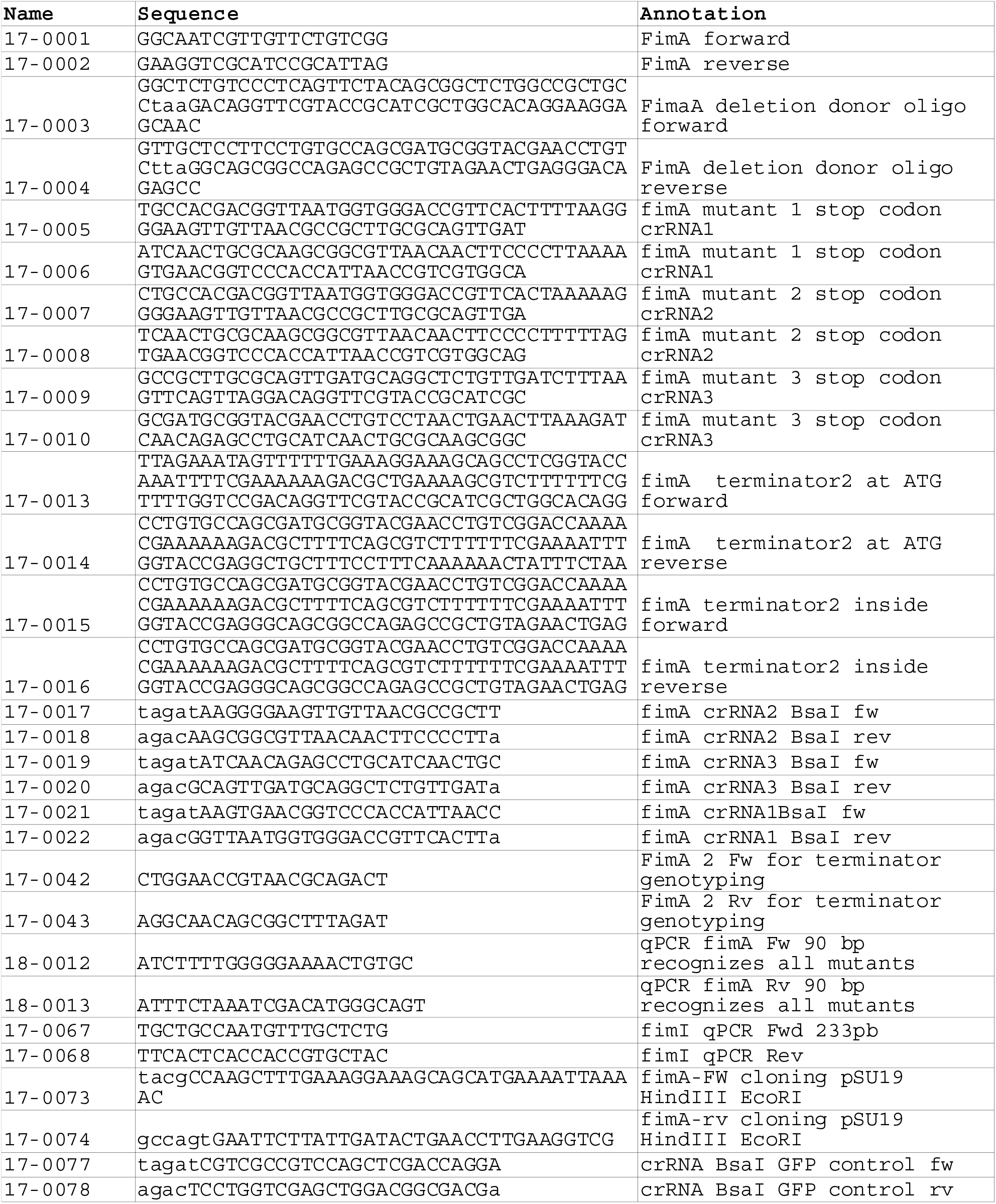
Oligos used in this study.

### AFM

To prepare AFM samples, punch-holed 1 cm pieces of muscovite mica were immobilized on metal supports using double sided tape. The static cultures were diluted ten times in ultrapure milliQ water. A drop of 100 μl from the diluted bacterial suspension was afterwards placed on the mica right after cleaving it with laboratory tape. The suspension was incubated for 20 minutes to allow adhesion of bacteria and the liquid was exchanged three times with ultrapure milli Q water. The samples were placed in the chemical hood and left to dry for a minimum of 5 min. The presence of bacteria on the mica was verified using an optical system mounted on the AFM (Bruker) and a random area was selected for imaging. The AFM imaging was performed in a MultiMode8 instrument (Bruker) in force tapping mode using the ScanAsyst software at a rate of 1 Hz and using ScanAsyst tips with a nominal spring constant of 0.4 N/m. Images shown in Fig. 6 depict the error signal generated by the piezo (J scanner).

## Acknowledgements

We thank the support of the Carl Tryggers Stiftelse för Vetenskaplig Forskning (projects CTS 15-96 and CTS18-65 awarded to David A. Cisneros), the Kempestiftelserna (projects JCK-1724 awarded to Bernt Eric Uhlin and SMK 1860 awarded to David A. Cisneros). We also want to thank Ramon Cervantes Rivera for helping with RNA purifications. We also thank Aster Assefa for her technical assistance in qPCR and the creation of deletion mutants and Andrea Puhar for critical reading.

---

**Supplementary Fig 1.**
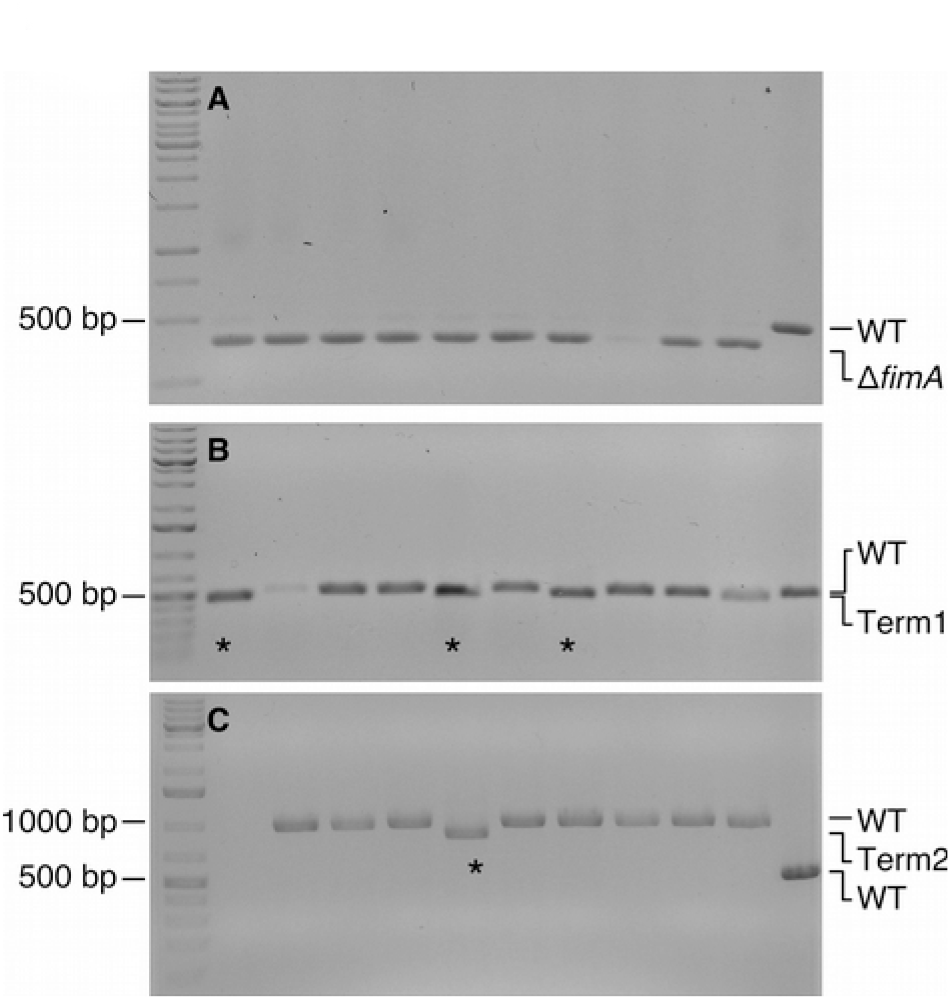
Efficiency of *fimA* mutant generation with Cas12a. A) 10/10 *E. coli* colonies co-transformed with crRNA1 and the donor oligo carrying a 97 bp *fimA* deletion tested positive for a mutation as detected by colony PCR. B) 3/10 *E. coli* colonies co-transformed with crRNA1 and the donor oligo carrying a 54 bp *fimA* terminator insertion tested positive as detected by colony PCR (marked with *). C) 1/10 *E. coli* colonies co-transformed with crRNA1 and the donor oligo carrying a 54 bp terminator insertion at the *fimA* start codon tested positive as detected by colony PCR (marked with *).

